# Interactive Neurorobotics: Behavioral and Neural Dynamics of Agent Interactions

**DOI:** 10.1101/2022.05.17.492233

**Authors:** Eric Leonardis, Leo Breston, Rhiannon Lucero-Moore, Leigh Sena, Raunit Kohli, Luisa Schuster, Lacha Barton-Gluzman, Laleh K. Quinn, Janet Wiles, Andrea Chiba

## Abstract

Interactive neurorobotics is a subfield which characterizes brain responses evoked during interaction with a robot, and their relationship with the behavioral responses. Gathering rich neural and behavioral data from humans or animals responding to agents can act as a scaffold for the design process of future social robots. The goals of this research can be broadly broken down into two categories. The first, seeks to directly study how organisms respond to artificial agents in contrast to biological or inanimate ones. The second, uses the novel affordances of the robotic platforms to investigate complex phenomena, such as responses to multisensory stimuli during minimally structured interactions, that would be difficult to capture with classical experimental setups. Here we argue that to realize the full potential of the approach, both goals must be integrated through methodological design that is informed by a deep understanding of the model system, as well as engineering and analytical considerations. We then propose a general framework for such experiments that emphasizes naturalistic interactions combined with multimodal observations and complementary analysis pipelines that are necessary to render a holistic picture of the data for the purpose of informing robotic design principles. Finally, we demonstrate this approach with an exemplar rat-robot social interaction task which included simultaneous multi-agent tracking and neural recordings.

## 1 Introduction

As technology and automation increasingly permeate every aspect of modern life, our relationship with these systems have begun to blur the lines between tool use and social interaction. Humans are routinely being asked to engage with artificial agents in the form of chat bots, recommender systems, and social robots which all exhibit facets of agency more familiarly associated with living beings (Saygin et al., 2012; Saygin & Cicekli, 2002; Gazzola et al., 2007). Such a promethean transition has raised an urgent need to study how biological organisms adapt and extend their social mechanisms to new digital simulacra (Baudrillard, 1981). The proliferation of technology has made sophisticated robotics accessible to many more scientists, empowering them to create a wide array of new applications, such as the use of robots to investigate animal models by physically interacting with those animals.

The goals of this research can be broadly broken down into two categories. The first, seeks to directly study how organisms respond to artificial agents in contrast to biological or inanimate ones. The second, uses the novel affordances of the robotic platforms to investigate coordination dynamics between agents Here we propose that to realize the full potential of the approach, both goals must be integrated as they provide complementary information necessary to contextualize one anothers results. For example, if one were to use a robot to study, without characterizing the organism’s differential response to robots and conspecifics, then it would be difficult to know whether the results reflected a generalizable reaction, or was an artifact of the animal’s particular response to the robot. Despite these epistemological liabilities, the degree to which animals’ responses differ between agent types, as well as the character of those differences, remains an open question.

At the nexus of our broadly construed categories is the nascent field, interactive biorobotics, that uses robots to experimentally probe questions in behavioral ethology, neuroscience, and psychology, using a variety of human and animal assays (Ishii et al., 2006; 2013; Gergely et al., 2016; Lakatos et al., 2014; Narins et al., 2003; Narins et al., 2005). Robot frogs that can emit auditory calls have been set up in environmental habitats and elicit fighting and even mating responses from wild frogs (Narins et al., 2003; Narins et al., 2005). Robot fish that interact with living schools of fish and vibrating robots that attract bees have shown effects on collective behavior in laboratory and naturalistic environments (Schmickl et al., 2021). Robot rats, like the Waseda Rat and the iRat, interact with living rats and affect their behavior in laboratory settings (Ishii et al., 2006; Wiles et al, 2012).

In this paper we provide strong evidence, using multimodal recordings from rats engaged in a social experiment with rats, robots, and objects. We show that interactions with agents of varying animacy and motion characteristics evoke different behavioral responses. In addition, we show measurable variations in the activity of multiple brain regions across different interactive contexts.

### 1.1 Background

Real organisms do not exist within a sterile world, acting against a quiescent backdrop. The world they adapted to is dynamic, inhabited by other agents, each behaving and interacting according to their own imperatives. Therefore, if science is to actually understand how the brain synthesizes stimuli and produces effective behavior it must tackle these complex, weakly constrained settings. It is technically complex to administer experimental manipulations in a minimally constrained environment, and the availability of new technology created an opportunity for experimentalists to partner with roboticists and computationalists to create data capture and analysis tools necessary to extract robust effects from the results.

Interactive biorobotics presents a promising new approach to facilitate experimental manipulation while retaining the complexities of inter-agent dynamics. As Datteri suggests, interactive biorobotics is a methodologically novel field that is distinct from classical biorobotics (Datteri, 2020). In classical biorobotics experiments, the robot is meant to “simulate” or replicate a function of a living system without direct interaction with the organism itself. In interactive biorobotics, the robot is meant to stimulate interaction with living systems and is instead bestowed with certain capacities to engage and stimulate a living system. The target of the explanation is the behavior of the living system in response to the robotic agent. A key aspect of interactive biorobotics is the integration and habituation of living systems to interactive robot counterparts (Quinn et al. 2018, Datteri, 2020). A popular example of an interactive biorobotics paradigm with animals was the Waseda Rat, a rat-like robot that continuously chases a living rat to induce anxiety-like behavior (Ishii et al, 2006; Shi et al, 2013).

Many experiments have demonstrated the viability of this technique for investigating neuroscientific questions. These results are the purview of interactive neurorobotics, a subfield which characterizes brain responses evoked during interaction with a robot, and their relationship with the behavioral responses. An early example of a pioneering interactive neurorobotics study is Saygin et al. (2012), which used neuroimaging to present human subjects with robots and androids for the purposes of studying the neural basis of the “uncanny valley.” This is an effect which shows that as a robot’s appearance increases in human-likeness, the more eerie or creepy it may seem to a human during interaction (Mori, 1970; Saygin et al, 2012). Behavioral evidence has been presented suggesting that macaque monkeys also have a similar response when presented with virtual avatars (Steckenfinger & Ghazanfar, 2009). However, it is unknown to what extent the uncanny valley effect is present in other animals, like rodents. While this is a fascinating question, this paper seeks to examine neural and behavioral responses to non-rodent-like robots with hopes of sidestepping any potential uncanny valleys leading to easier acceptance.

An example of an interactive neurorobotics experiment, where the brain is measured along with animal behavior, was conducted with the predator-like robot known as “Robogator.’’This robot alligator that can walk and bite, interacted with rats in a dynamic foraging task (Choi & Kim, 2010). In their task, as a rat approaches a food reward, the Robogator suddenly snaps its jaws towards the rodent, resulting in an approach-avoidance conflict paradigm. In this experiment, Choi and Kim (2010) drastically inhibited amygdala function by locally infusing muscimol (a GABA agonist) and inducing electrolytic lesions. They found that without amygdala function, animals showed diminished fear responses towards the robot. They also demonstrated that local infusions of muscimol globally suppressed amygdala activity, leading to increased exploration of the robot. Similar approach and avoidance dynamics are present in a variety of behavioral paradigms related to predation, social interaction, reward learning, and threat detection (Jacinto et al., 2016; Mobbs & Kim, 2015). The increase in exploration under conditions in which the amygdala is suppressed underscores the importance of the state of the animal in approaching potentially threatening, frightening, or unknown stimuli.

When potential danger or even conditions of high uncertainty are present, there is a cost to exploration due to rats’ natural propensity for neophobia (Mitchell, 1976; Modlinska et al., 2015). Inherent in introducing robot counterparts to rats is not only the uncertainty of novelty but also risk assessment regarding the fear of harm. Thus, in order to explore robots, rats must engage in regulatory behaviors allowing the neural system to enter states that allow exploratory behaviors. Inevitably, this necessitates a balance between the sympathetic nervous system and the parasympathetic nervous system, invoking allostatic processing in which systemic stability is achieved through continuous change. Robots have been more often used in fear and stress inducing paradigms with novel robots that exert extreme regulatory demands on the rat (Ishi et al, 2006; Choi and Kim, 2010). During the management of uncertainty, self-regulatory behaviors actively modulate internal demands to meet external demands (Nancy & Hoy, 1996).

In addition to examining behaviors related to novelty and fear, studies have also been performed examining the similarity between rodent social behaviors and behaviors toward a robot. Rats have been shown to behave similarly towards mobile robots as they do to rats, exhibiting behaviors such as: approaching, avoiding, sniffing, and following (del Angel Ortiz et al, 2016; Wiles et al, 2012; Heath et al, 2018). The analysis showed that the rats demonstrated similar relative orientation formations when interacting with another rat or moving robot. Analysis of relative spatial position in rat-rat dyads in comparison with rat-robot dyads show similarities, raising the question of whether these dynamic interactions might have social elements (del Angel Ortiz et al, 2016). Endowing robots with coordination dynamics that mitigate concerns of dominance still leave uncertainty regarding potential animacy of the agent.

A robot’s ability to engage in self-propelled motion is crucial for its potential to elicit social and attentive behaviors. Studies of social recognition and novel object recognition traditionally disregard dynamic objects that exhibit self-propelled movement despite early demonstrations that even two-dimensional objects that exhibit self-propelled movement are often associated with sociality, agency, and animacy detection (Heider & Simmel, 1944). Self-propelled motion can take various forms. Biological motion is associated with maximizing smoothness and can be recognized as animate even with minimal representation (Flash & Hogan, 1985; Todorov & Jordan, 1998; Saygin et al., 2004). Mechanical and materials constraints on interactive robots make the use of biological motion rare and difficult to achieve. However, interactive robots provide an opportunity to experimentally manipulate agency by utilizing elements of animation through movement trajectories and temporal coordination (Hoffman & Ju, 2014).

A robotic platform that displayed emergent semi-naturalistic movement patterns is the iRat neurorobotics platform (Ball et al., 2010, Wiles et al, 2012). The iRat was originally developed with a neurally feasible model that simulated the function of the hippocampus, entorhinal cortex and parietal cortex for the purpose of learning spatial environments (Ball et al., 2010). By virtue of entering an environment and exhibiting appropriate spatial navigation and object avoidance, iRat garnered observational attention from on-looking rats (Wiles et al.,2012). This brought about the question of whether iRat could be driven to behave in a socially interactive manner that would result in rats engaging in prosocial behavior with the iRat as they do with conspecifics (Rutte & Taborsky, 2008; Bartal et al., 2011). Quinn et al. (2018) used the iRat to interact with rats for the purpose of eliciting social responses and examining prosocial behavior towards robots (For Artists Depiction and Picture of Live Interaction See Figure 1). Rats will not only engage in prosocial behaviors with each other, such as freeing other rats from an enclosed restrainer, but have also been shown to reciprocate with robots (Quinn, et al. 2018, Wiles et al., 2012).

**Figure 1.**
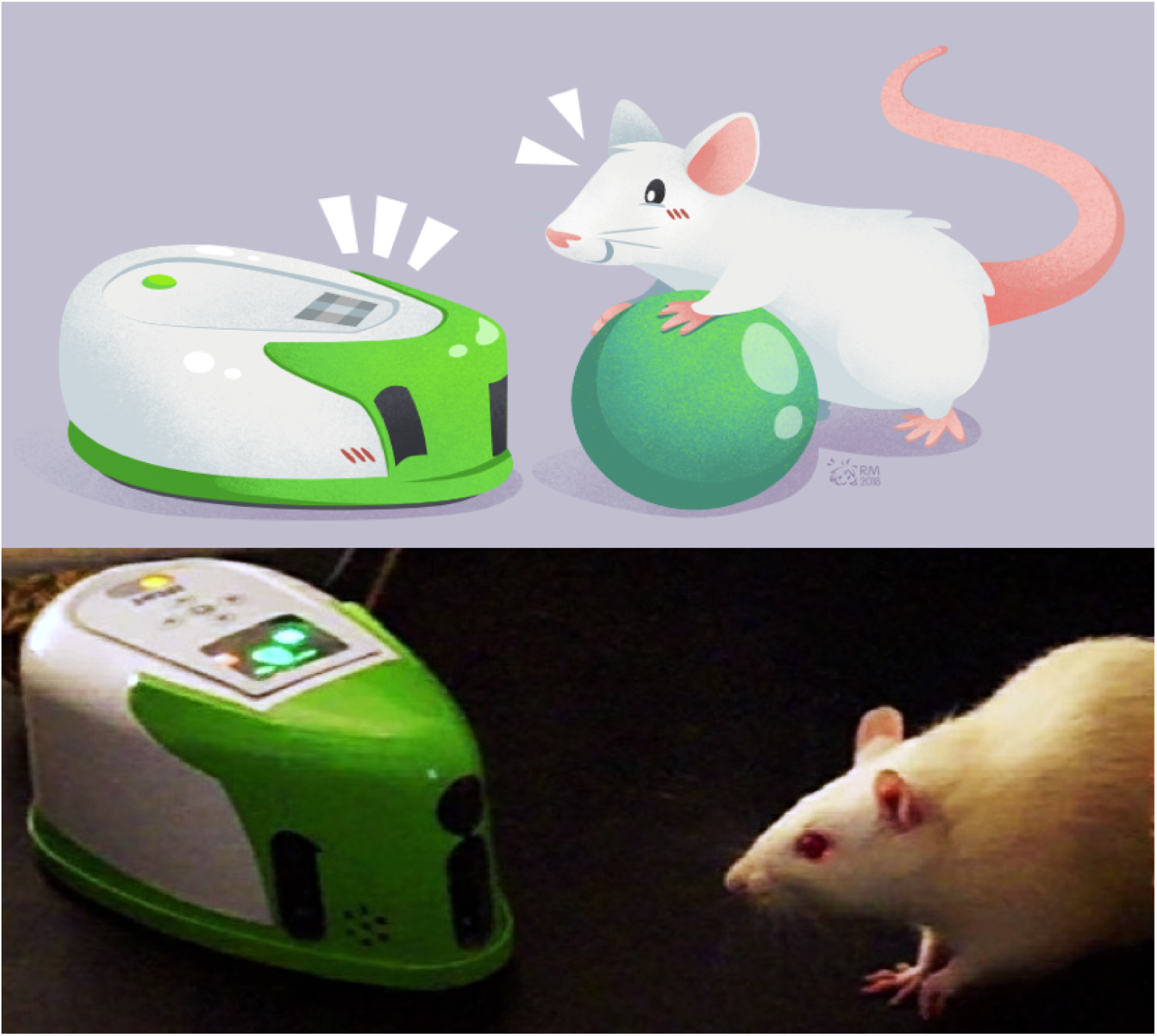
A, An artist’s depiction of rat-robot and rat-object interactions from Quinn et al., 2018 (photo credit pending permission from Rosana Margarida). B, A picture of live rat-robot interaction.

In this paper we further prior work by investigating the effect of agent type on both behavioral and neural outcomes. For our experimental model we considered rats engaging in a socio-robotic experiment developed according to principles of rodent-centered design (See Figure 1). In this study, rats interacted in a circular arena with other rats, robots, or objects. They engaged in naturalistic behaviors during this time and the resultant observational data was used to characterize the dynamics of rats’ behavioral repertoires in the presence of different agents or objects. While the rats were freely behaving, we used multi-site electrophysiological recordings to examine brain states during different behavioral states. We examine how the agent-based interactions influence neural oscillations within local field potentials in the olfactory bulb, amygdala and hippocampus (see Brain Areas of Interest) during grooming, immobility, and rearing behaviors (see Behaviors of Interest). To investigate dynamic interactions, we demonstrate the use of deep learning video tracking for offline multi-animal and robot tracking.

### 1.2 Brain Areas of Interest

The neural circuit examined in this paper was chosen because it spans specific functional aspects critical for social behavior. Together, the olfactory bulb, amygdala, and hippocampus form a tightly connected network (for review see Brodal, 1947) that provide information about autonomic, sensory, spatio-temporal, and affective context (Moberly et al, 2018, Jacobs, 2022) (See Figure 2). They are, thus, well situated to provide valuable information regarding the benefits and liabilities of the external environment during social interaction, exploration, and sampling of the sensory information. Further, they have well characterized anatomical connectivity and diverse observed forms of frequency dynamics which makes them an excellent subject for studying the holistic closed loop processing of social information.

**Figure 2.**
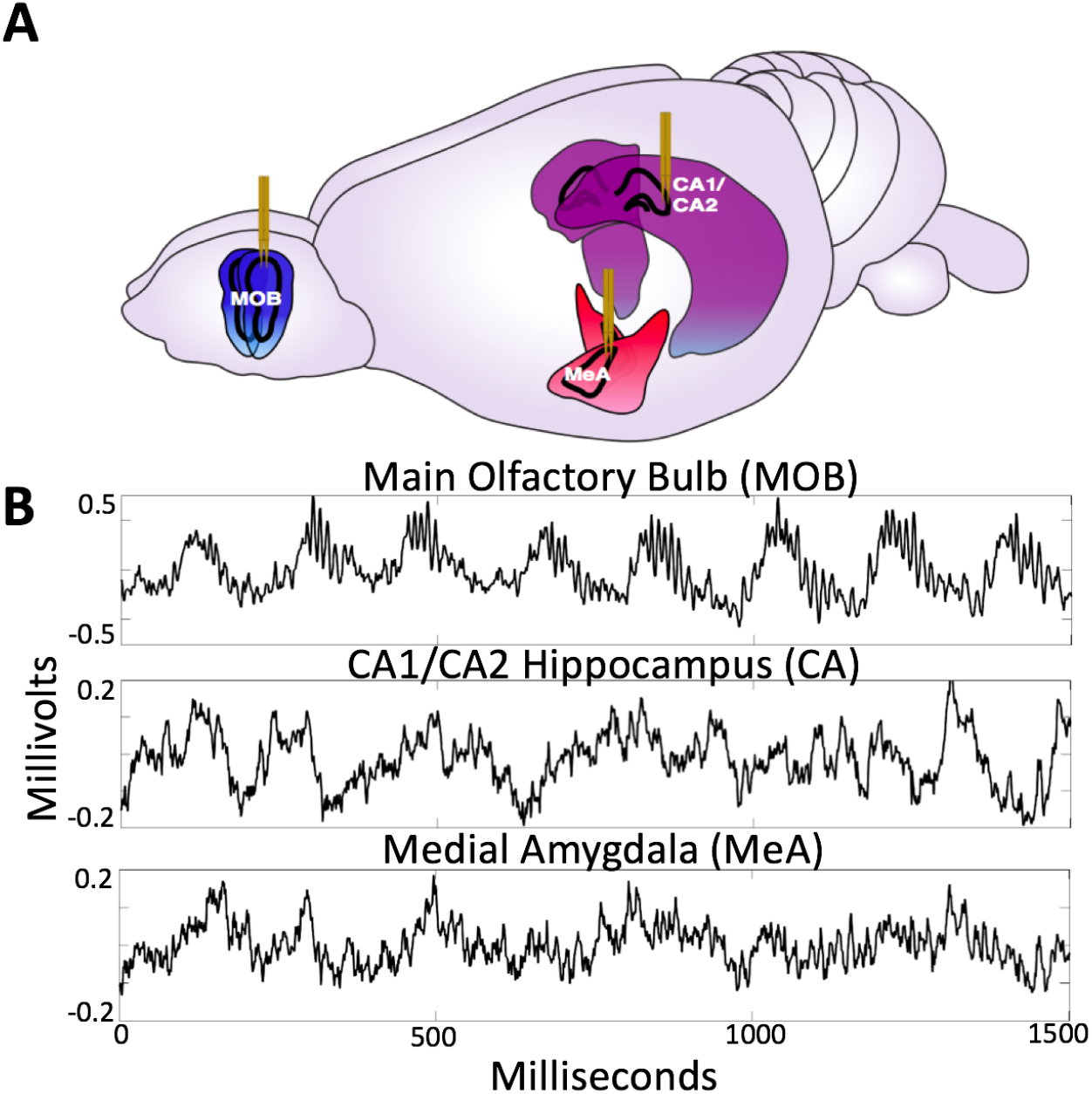
A, Diagram of electrode placement in the Main Olfactory Bulb (MOB), Medial Amygdala (meA), Hippocampus (CA1/CA2). Figure adapted from scidraw.io under Creative Commons 4.0 license (Tang, 2019). B, Raw traces from the MOB, Ca1/Ca2 and meA.

The **olfactory bulb** is a dominant primary sensory organ for rodents, playing a central role in odor discrimination and social cognition for rats (Dantzer et al., 1990). The main olfactory bulb (MOB) local field potential exhibits neural oscillations associated with sensory processing (Kay et al., 2009). Kay et al. (2014) have shown that the frequency of the theta oscillation in the olfactory bulb LFP reliably follows respiration rate (2-12Hz). Respiration-entrained oscillations provide information about the state of the autonomic nervous system, and are distinct from theta oscillations (Tort et al., 2018). It is important to note that respiratory rhythms are not restricted to the olfactory bulb, and are also found throughout the brain (Heck et al., 2017; Rojas-Libano et al., 2014). The following study seeks to highlight the important contribution of olfaction and respiratory-related brain rhythms to brain dynamics (Jacobs, 2012; Lebedev et al., 2018), in order to simultaneously examine sensory and autonomic dynamics as they relate to agent and object interactions. Olfactory and hippocampal theta oscillations also show coherence during olfactory discrimination tasks (Kay, 2014). Recent work also suggests coupling of the beta rhythm from olfactory bulb to hippocampus, which suggests a directionality of functional connectivity going from OB to hippocampus (Gourévitch et al., 2010). The olfactory bulb also exhibits two distinct gamma oscillations associated with contextual odor recognition processing, low gamma (50-60Hz) during states of grooming and immobility and high gamma (70-100Hz) associated with odor sensory processing (Kay, 2005). Future work will highlight the role of these rhythms, whereas the current work focuses on dynamics within the theta, respiratory, and beta frequencies. (For raw traces of the MOB LFP see Figure 2B).

The **amygdala** is a complex of historically grouped nuclei located in the medial temporal lobe, commonly associated with affective processing, saliency, associative learning, and aspects of value (Gallagher and Chiba, 1996). Oscillations in the amygdala and their coherence with other brain structures have been linked to learning and memory performance (Paré et al., 2002). The function of the medial amygdala (MeA) is associated with the accessory olfactory system, which receives inputs from the vomeronasal organ, playing a role in social recognition memory, predatory recognition, and sexual behavior (Bergan et al., 2014). The MeA receives input from accessory olfactory areas and has projections to hypothalamic nuclei that regulate defensive and reproductive behavior (Swanson & Petrovich, 1998). The basomedial nucleus has been linked to the regulation and control of fear and anxiety-related behaviors (Adhikari et al, 2015). Amir et al (2015) investigated how principal cells in the basolateral amygdala respond to the Robogator robot during a foraging task. A group of cells reduced their firing rate during the initiation of foraging while another group increased firing rate. It was found that this depended on whether the rat initiated movement, with the authors’ suggesting that the amygdala is not only coding threats and rewards, but also is closely related to the behavioral output. (For raw traces of the MeA LFP see Figure 2B).

The **CA1/CA2** subregion of the **hippocampus** is functionally associated with spatial navigation, contextual information, and episodic memory formation. The CA1 region of the hippocampus is commonly associated with spatial navigation dorsally and social/affective memory ventrally (van Strein et al., 2009). The hippocampal theta rhythm is a 4-10Hz oscillation generated from the septo-hippocampal interactions, which temporally organize the activity of CA1 place cells according to the theta phase. The CA2 region of the hippocampus plays an important role in modulating the hippocampal theta rhythm and also an essential part in social memory (Mercer et al., 2007; Hitti & Siegelbaum, 2014, Smith et al 2016). CA2 receives projections from the basal nucleus of the amygdala, which plays an important role for contextual fear conditioning (Pitkanen et al., 2000; Goosens & Maren, 2001). Ahuja et al. (2020) demonstrated that CA1 pyramidal cells were responsive to a robot that indicates a shock zone. Robots have also been developed for the purpose of improving behavioral reproducibility when examining aspects of rodent spatial cognition in neuroscience. When rats navigate through a maze, there can often be a variety of factors that an experimenter might want more control over. In this case a robot with an onboard pellet dispenser was used to regulate the rat’s direction and speed as they moved through the maze (Gianelli et al.,2018). (For raw traces of the CA1/CA2 LFP see Figure 2B).

### 1.3 Behaviors of Interest

The types of behavioral epochs used for the comparative neural analysis were chosen due to their ready identifiability and representation of distinct types of exploratory and self regulatory behavior. The behaviors are generally demonstrated when the rat temporarily stopped running or walking and was not physically exploring the other agent on the open field. The behaviors we prioritized include immobility, self-grooming, and rearing (See Figure 3). Given that the state of the rat differed widely across behaviors, it was particularly important to examine how the presence of different agents (rat, robot, or object) perturbed their state within a particular behavior.

**Figure 3.**
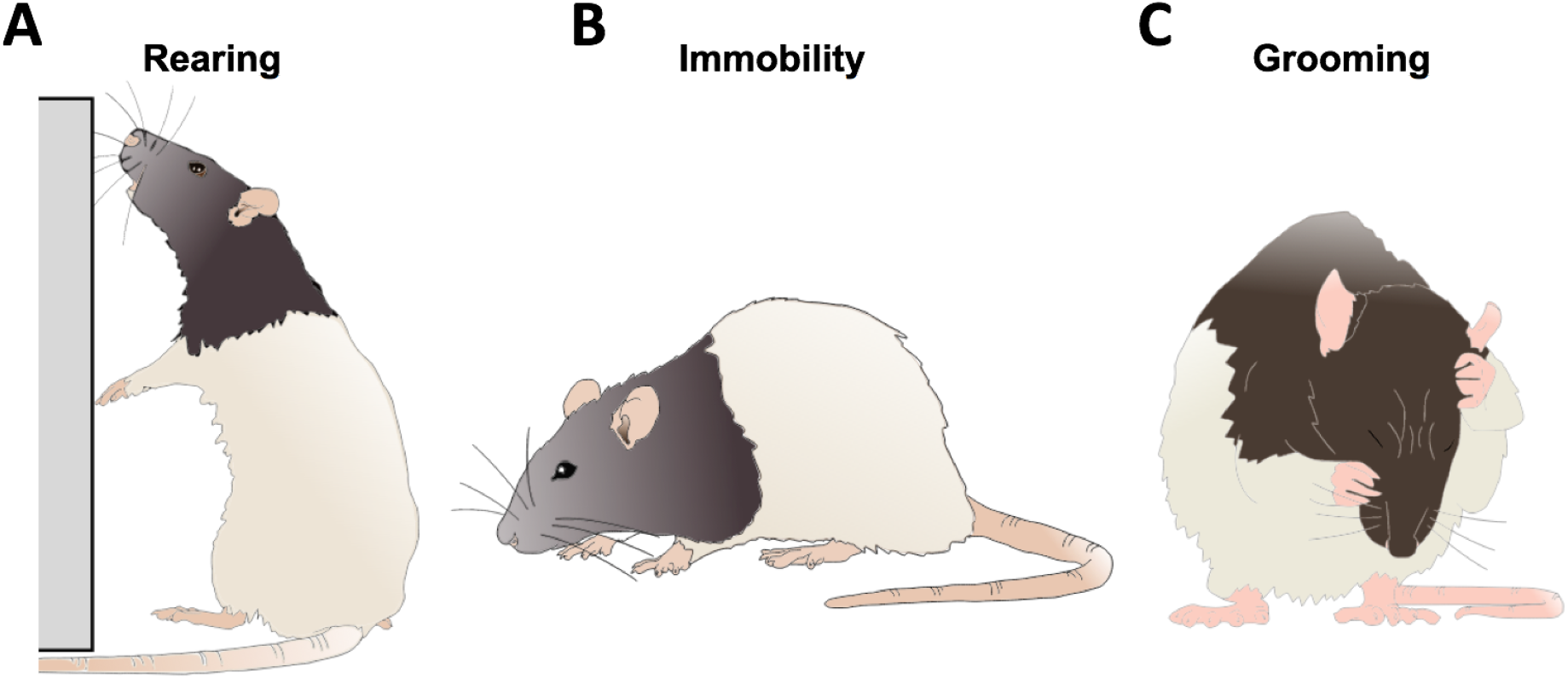
A depiction of a rat engaging in rearing, immobility and grooming behaviors. Figure adapted from scidraw.io under Creative Commons 4.0 license.

**Immobility** is a complex behavior distinguished by the motionlessness exhibited by the subject. This behavior is made distinct and significant as an intermediary behavior between the regulatory and exploratory states rats can alternate between. During immobility, the rat is likely in a heightened state of arousal, with an intensity of alertness. With current comprehension, engaging in immobility is a behavior indicative of risk assessment, occurring preemptively to a threat or as a result of one (Kay, 2005). Immobility allows the rat time to acquire and interpret environmental stimuli, triangulate any potential discomfort or stressors, and act accordingly.

**Self-grooming**, hereafter referred to as grooming, is a behavior inherent to rodents that is communicative of not only hygienic regulation, but also self-regulation of stress relief (Fernandez-Teruel & Estanislau, 2016). Grooming is a regulatory process, which often serves the function of de-arousal (Kalueff et al., 2016). Grooming includes sequences of rapid elliptical strokes, unilateral strokes, and licking of the body or anogenital area. Due to the behavioral complexity of grooming, frequency and duration of grooming bouts is dependent on context (Song, Berridge, & Kalueff, 2016).

**Rearing** is an exploratory action, exhibited as a means of increasing the rat’s access to stimuli in the environment. In both instances, the rat straightens and lengthens their spine and maneuvers their forelegs to increase their height. When rearing, rats increase their sensory exploration of the surrounding environment (Lever, Burton, & O’Keefe, 2006). This fluid behavior varies greatly in its duration and frequency. For the purposes of this paper, rearing was specifically defined with relation to the wall of the environment.

## 2 Materials and Methods

### 2.1 Animals and Housing

All experiments and maintenance procedures were performed in an American Association for Accreditation of Laboratory Animal Care (AAALAC) accredited facility in accordance with NIH and Institutional Animal Care and Use Committee (IACUC) ethical guidelines and preapproved by the IACUC committee. 6 Sprague-Dawley rats (n = 6) (Harlan Laboratories) performed in the behavioral experiments. 3 rats (n = 3) were surgically implanted with electrodes for electrophysiological recordings. They were acquired at 6 weeks old and housed in pairs. Cagemates were put together in an enriched environment for 30 minutes a day and were maintained on a 12 hour day/night cycle. After receiving surgery, the implanted rat was single-housed for the whole of the experiment. To offset the lack of social enrichment from being single-housed, they were taken out to play in the enriched environment with the former cagemate on the same schedule.

### 2.2 Experimental Conditions and Trial Information

Trials were collected from a rat and a robot freely roaming in an arena (n = 40). Comparison conditions included object trials (n = 21), social interaction trials with other rats (n = 20), and solo open field exploration (n = 84) (See SI for Animals and Housing). Trial lengths were approximately 3 minutes long. Trials were counterbalanced for order effects. The robots and the animals’ were recorded using an overhead camera. The videos were used to hand-label behaviors and estimate position of the interacting agents for each video frame (See SI Behavioral Coding). Rats were surgically implanted with electrodes in the main olfactory bulb, hippocampus and amygdala simultaneously to record local field potentials during freely moving behavior (See SI for Surgical Procedure and Neural Implants and Recordings).

### 2.3 Robot

The iRat (n = 2) is a robotics and modeling platform created by the Complex and Intelligent Systems Laboratory (Ball et al, 2010). iRat is a two wheeled mobile robot that is 180mm x 100 mm x 70 mm. The iRat is capable of both WoZ interaction and performing pre-programmed behaviors. Robots were distinguished visually by color (red and white iRat / green and white iRat) and using distinguishing olfactory odors. The Red iRat was tagged with frankincense essential oil, the green iRat was tagged with myrrh essential oil. These odors were chosen in order to match preference profiles and are within the same category of woody scents. Our lab has previously shown that rats do not demonstrate a preference for either scent (Quinn et al, 2018). For information about the experimenter’s control of the robot’s movement dynamics see SI Wizard of Oz (WoZ). For the experiment, locomotion of both robots was limited and reduced to below 0.5m/s. In addition to the iRats, multiple Arduino-based mobile robots were also used (see Arduino Board of Education Shield Robots in the Supplemental Information).

### 2.4 Automated Tracking-Based Behavior Segmentation

Let S refer to the agent’s state which is the position in Cartesian x,y coordinates over time and θ orientation in radians over time t by video frames (See Figure 4). The position and orientation for the rat is denoted by *S_r_* = {*x_rat_*, *y_rat_*, θ_*rat*_} and the robot *S_ρ_* = {*x_robot_*, *y_robot_*, θ_*robot*_,}. Inter-agent distance (IAD) was calculated by taking the euclidean distance between *S_r_* and *S_ρ_* position vectors.

**Figure 4.**
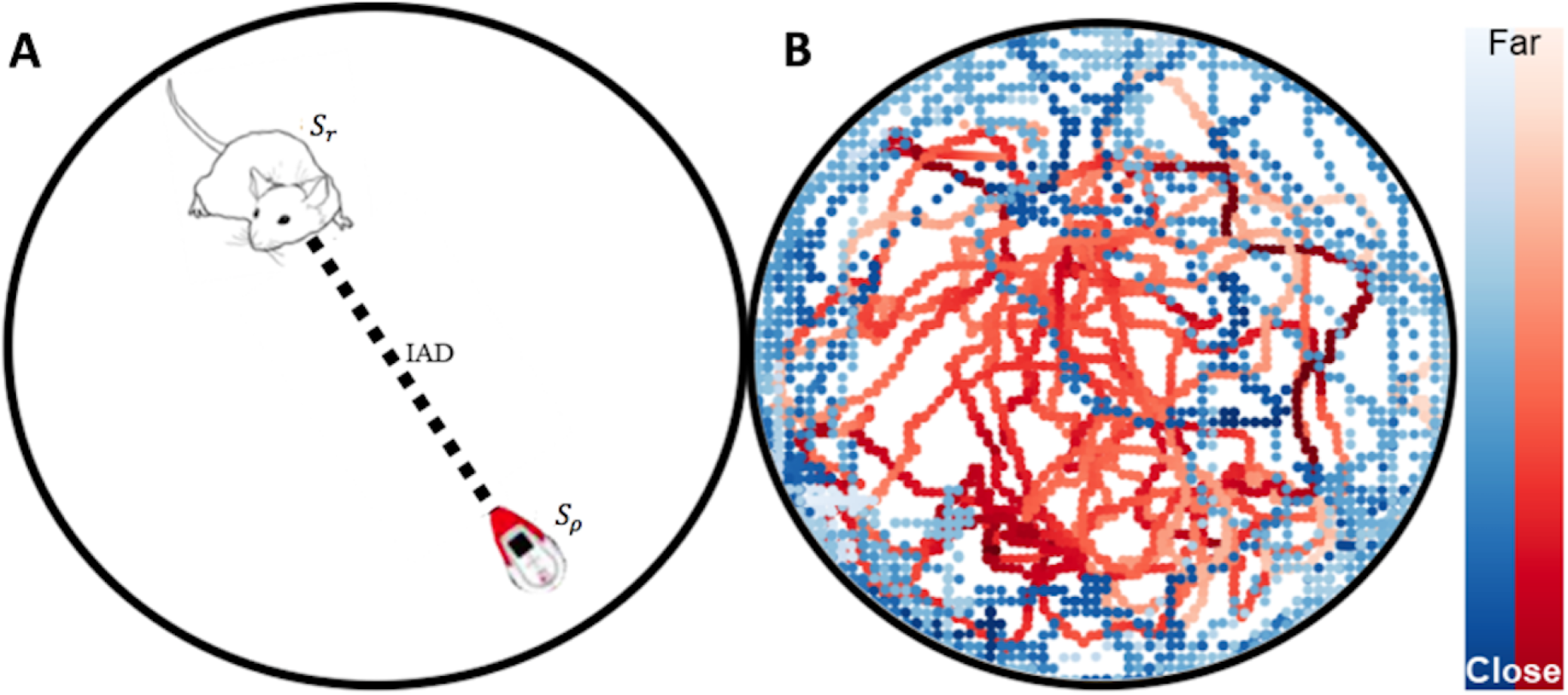
A, Animal and Robot Position Tracking With SLEAP. Automated tracking-based behavior segmentation with state vectors *S_r_* and *S_ρ_*, and the inter-agent distance (IAD). B, Tracking data from a robot freeroam trial is plotted.

### 2.5 Behavioral and Tracking Statistics

For the tracking data, inter-agent distances were calculated using the euclidean distance between agents for the rat-rat, rat-robot and rat-object interaction conditions. The distributions of inter-agent distances, mean event counts per trial, and mean event duration per condition were compared using one way Welch’s t-tests, and effect sizes were calculated using Cohen’s d. For the behavioral events, the event frequency was calculated per trial and the mean duration for each behavioral event type was calculated per condition. The events and durations were also compared using one way Welch’s t-tests, and effect sizes were calculated using Cohen’s d.

### 2.6 Mixed Effects Model

A general linear mixed-effect model was constructed to perform an ordinary least squares regression of a response variable as a function of mixture of fixed and random effects. Fixed effects include the behavior and agent type, while random effects include the influence of the variance of each individual rat on the response variable within the region. Null distributions were created by taking the aggregate average of all behavioral events. This allows for the comparison of changes in average power within rats, while controlling for any uneven sample sizes and individual differences in overall power within brain regions. The effect size was estimated by subtracting the means in question and dividing by the standard deviation of the residual. The mean coefficients, standard errors, z scores, p values, and effect size estimates are reported. The intercept of the baseline group is reported as Int., standard error of the mean as SEM, and the mean coefficients of the comparisons are reported as M. Z scores and p values are also reported.

## 3 Results

### 3.1 Tracking Results

The inter-agent distances maintained during interaction were minimal between rats, slightly longer for rats and objects, and the longest for rats and robots (See Figure 5). The inter-agent distances for rat-robot interactions indicate that the rat and robot maintain a significantly longer distance on average (M = 167.09 pixels, SEM =.14) than interactions with conspecifics (M =110.20 pixels, SEM = 0.21, t =227.86 pixels, p < e-10, d = .75). The rat-robot inter-agent distance distribution had a significantly higher mean than the rat-object distance distribution (m=131.44, sem=.22, t=135.56, p < e-10, d = .48) The rat-object inter-agent distance distribution showed a significantly higher mean than rat-rat distances (t = 69.93, p < e-10, d = .28).

**Figure 5.**
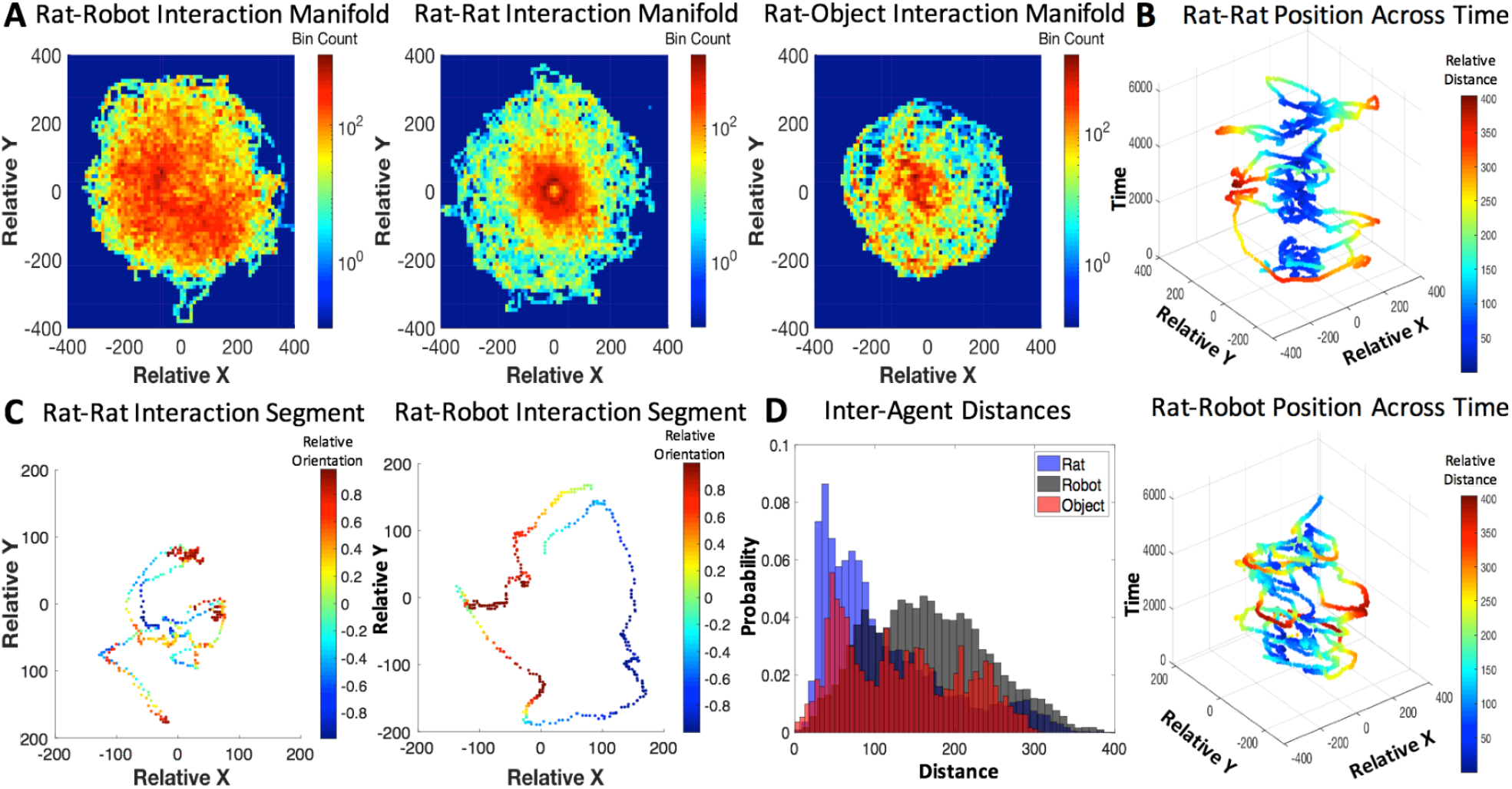
A, Agent-based interaction manifolds represented by 2D histogram bins along the relative x and y position axes. B, Relative position over time for the rat-rat and rat-robot interactions with a colormap revealing inter-distance between agents C, shorter segments of relative position of rat-rat and rat-robot interactions with a colormap revealing relative orientation. D, a histogram of inter-agent distance per rat, robot and object conditions.

### 3.2 Behavioral Hypothesis Tests

For a summary of the hand-coded behavioral event identification please see SI Behavioral Video Coding.

#### 3.2.1 Immobility

It was predicted that interaction with the robot leads to increased risk assessment behaviors, signified by increased immobility behavior (See Figure 6). The mean frequency of immobility events during rat-robot interactions (M = 2.70, SEM = .37) per trial was significantly larger than those from rat-rat interactions (M = 1.31, SEM = .31, t = 3.29, p = .001, d = .75). The mean duration of immobility during open field interactions (M = 4.95, SEM = .30) was significantly longer than robot (t = 2.73, p < e-4, d =1.44), rat (t = 8.77, p < e-4, d = 1.15) and object (t = 9.06, p < e-4, d = 1.40) interactions.

**Figure 6.**
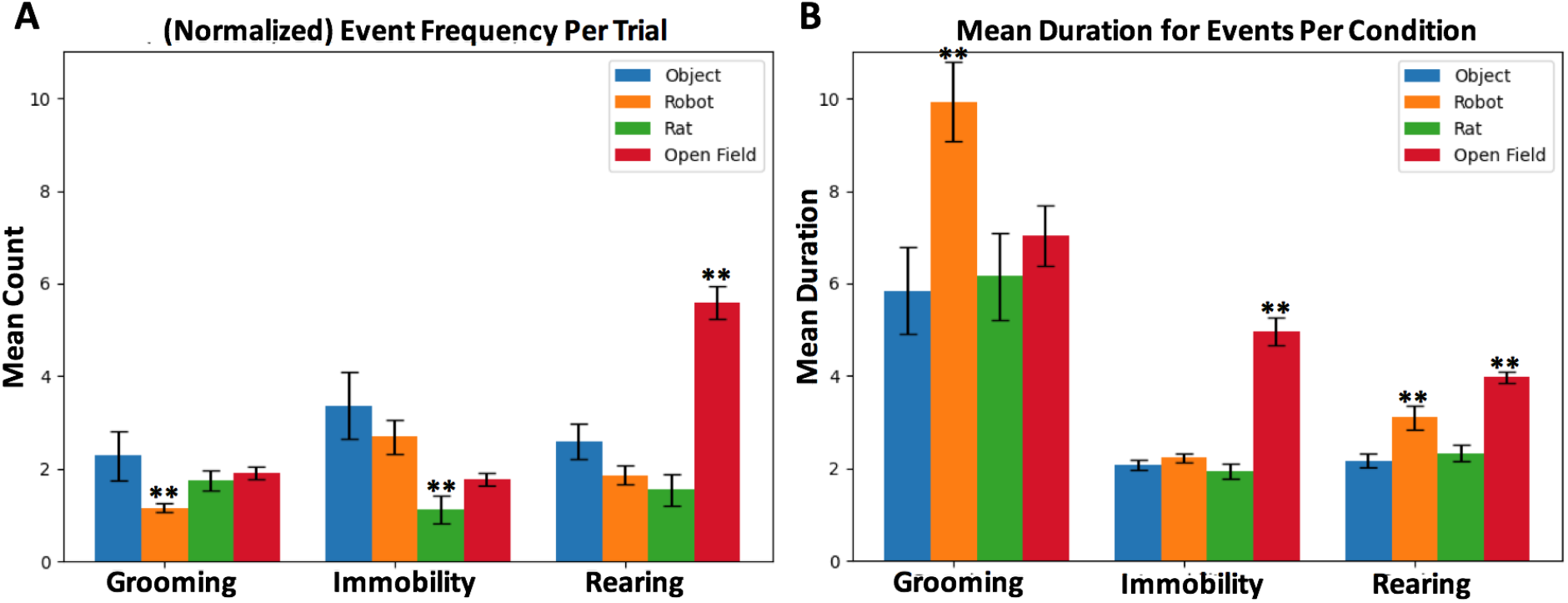
Mean counts per trial and mean duration per condition for grooming, rearing and immobility events. Error bars indicate the standard error of the mean.

#### 3.2.2 Grooming

Another hypothesis was that the robot would perturb grooming behavior due to distress related to a possible threat and stimulus uncertainty (See Figure 6). During interaction with the robot, although grooming events were significantly less frequent (which was counter to our prediction) on average per trial (M = 1.16, SEM = .11) than the rat (M = 1.74, SEM = .22, t = 2.38, p = .012, d = .82) and object interactions (M = 2.29, SEM = .11, t = 2.07, p < .04, d = 1.00). Despite being less frequent, grooming events during interaction with the robot (M = 9.93, SEM = .85) show a marginally longer duration than interaction with rats (M = 6.15, SEM = .94, t = 1.78, p < .04, Cohen’s d = .36) and objects (M = 5.8, SEM = .93, t = 1.93, p < .03, d = .39), indicating an alteration in duration of distress related grooming in the presence of robots.

#### 3.2.3 Rearing

It was predicted that the robot and open field will result in increased rearing as escape-related exploratory response (See Figure 6). Rearing events during interaction with the robot (M = 3.09, SEM = .26) show a significantly longer duration than rat (M = 2.32, SEM = .18, t = 2.36, p = < .01, Cohen’s d = .30) and object (M = 2.17, SEM = .16, t = 2.94, p < .002, d = .38), confirming the hypothesis that the presence of a robot might elicit exploration. The mean duration of rearing events during open field interactions (M = 3.9, SEM = .12) was significantly longer than the robot (t = 2.99, p < .002, d = .34), object (t = 8.99, p < e-4, d = .76), and rat (t = 7.62, p < e-4, d = .69) conditions. Empty open fields typically elicits exploratory vigilance, thus it appears that the open field induces an increase in hypervigilance relative to the robot.

### 3.3 Neurophysiological Results

Neurophysiological signals were recorded in rats during their behavioral displays (grooming, immobility, and rearing) occurring throughout interaction sessions with different agents (rat, robot, object). The behavioral epochs were extracted from continuous data and occur naturally throughout the interaction sessions. This means that the display of behaviors varies both with respect to inter-agent position and order of occurrence. The event durations of these naturalistic behavioral epochs differ, so as not to impose artificial cut-offs on natural display of behaviors.

#### 3.3.1 Spectrograms

To demonstrate an example of inter-agent variation within one behavior type, spectrograms below (see Figure 7) show multi-region brain dynamics from periods in which implanted rats exhibited immobility behaviors during rat, robot or object interactions. The pre- and post-event windows are 1 second in length providing a sense of scale for the viewer. In Panel A the period of immobility to a rat is approximately one second and in Panel B the period of immobility to robot is approximately 3 seconds.

**Figure 7.**
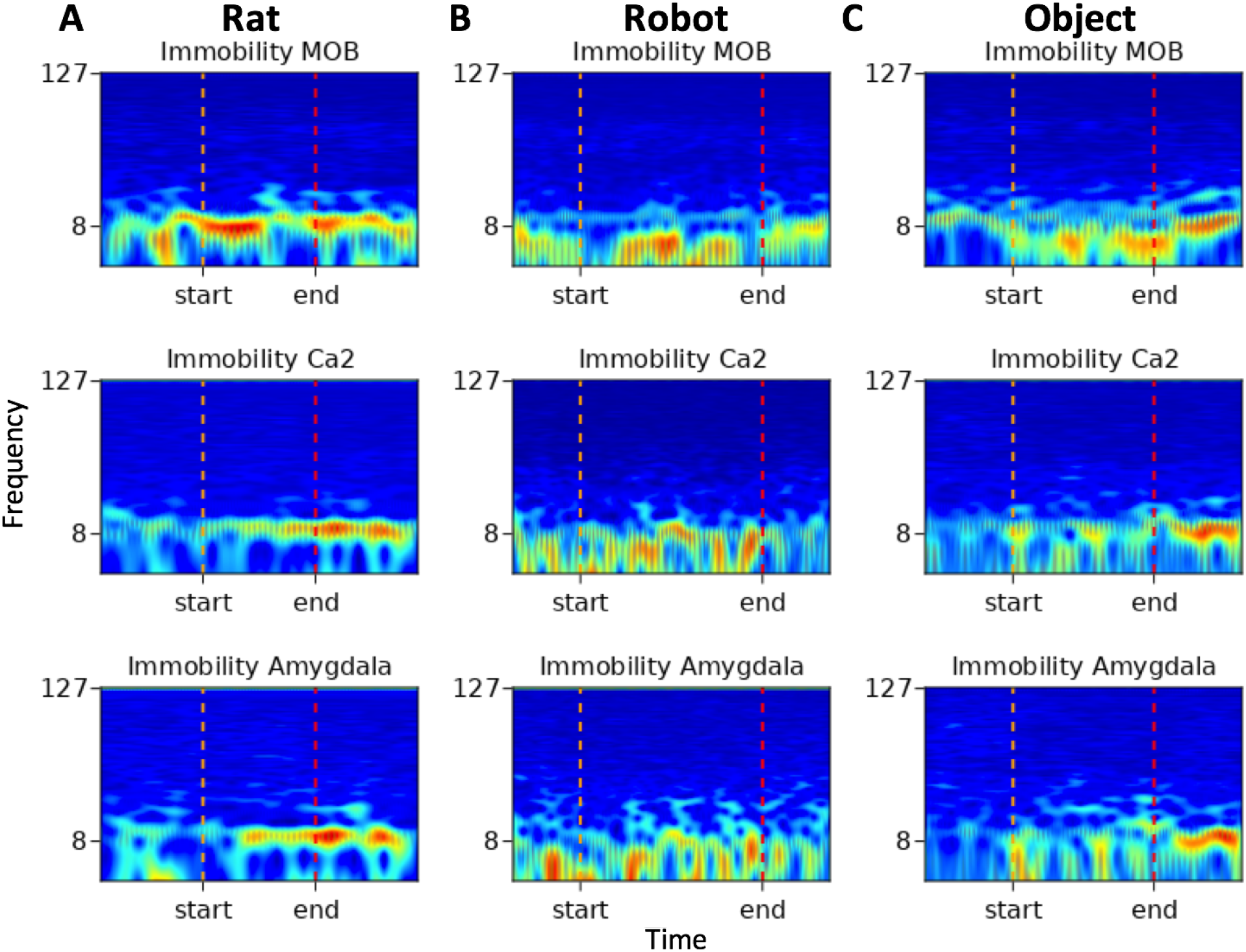
Spectrograms of immobility events from rat, robot and object conditions. Frequency (Hz) is represented along the y-axis with markers for 8Hz and 127 Hz, time is represented along the x-axis, the colormap represents high amplitude in red and low amplitude in blue. The start and end indicators denote the onset and offset of the variable duration behavioral events.

The immobility event which occurred during rat-rat social interactions in Figure 7 Panel A shows increased amplitude within the theta range in all brain regions with some transient beta oscillations in MOB and amygdala towards the end of the event. The immobility events that occurred during robot and object interactions in Figure 7 Panels B and C show high amplitude in respiratory oscillations in all regions with more pronounced beta oscillations in the amygdala during the event.

#### 3.3.2 Power Spectral Densities

### 3.4 Neural Hypothesis Testing

The results of the mixed effects model showed a variety of effects in the respiratory, theta, and beta frequency ranges (See Figure 8 for average power spectral density plots). There are also marginal effects in the gamma range that will require a larger data set for validation and will not be addressed in this study.

**Figure 8.**
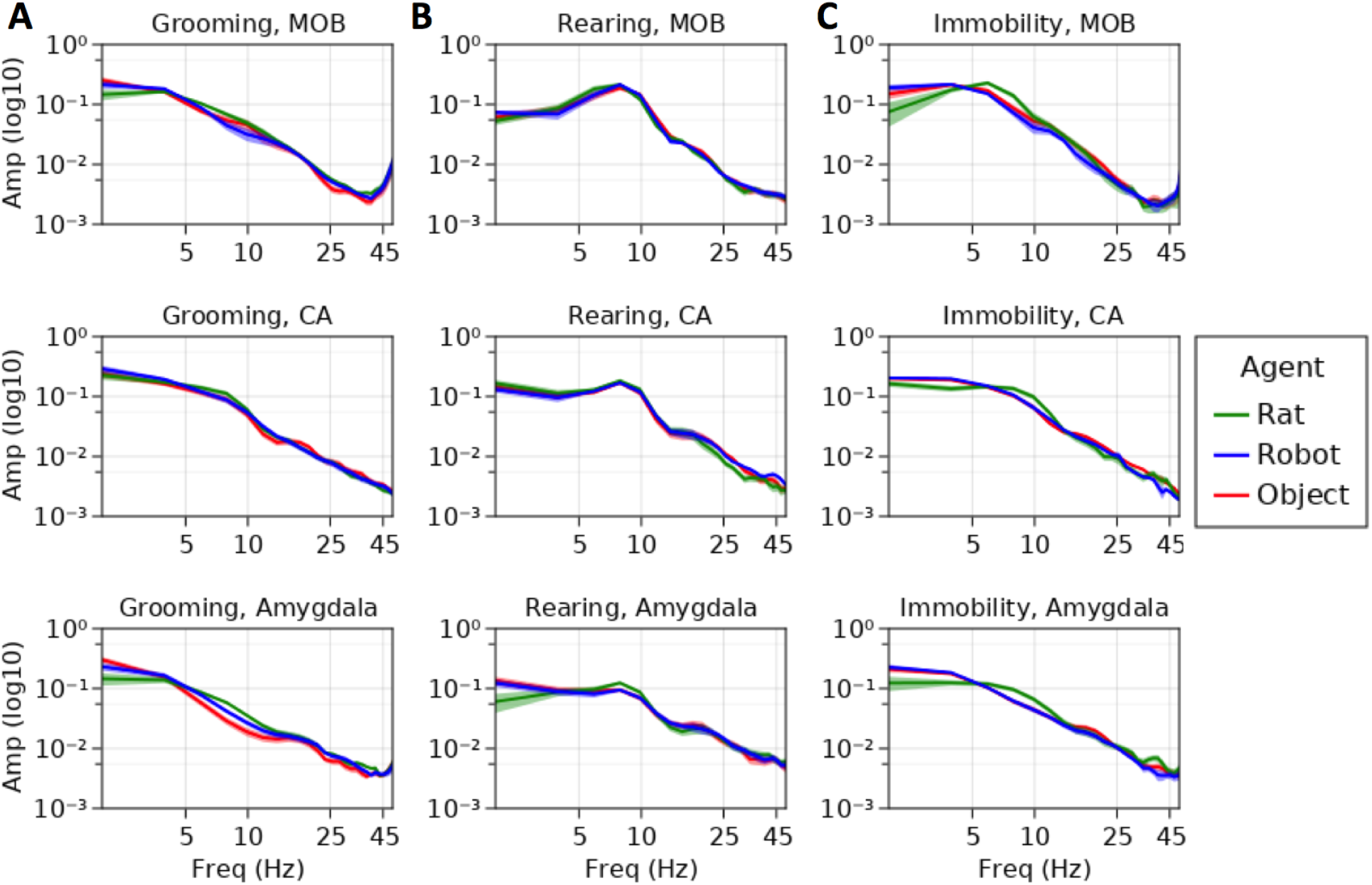
Average power spectral densities for MOB, CA1/CA2, and Amygdala for A, grooming, B, rearing, and C, immobility events. Amplitude is represented on the y-axis, frequency is represented along the x-axis. Average amplitudes during interaction with rats is represented in green, robots in blue and objects in red.

#### 3.4.1 Theta Oscillations: Olfactory Exploration and Salience

Other conspecifics were expected to elicit heightened sensorimotor exploration relative to robots and objects (See Figure 8). Rats are inherently more complex olfactory stimuli, and this is indicated by theta oscillations. Immobility events showed significantly higher theta amplitude in MOB during the rat-rat interaction trials (Int = .026, SEM = .0055) compared to the events from the robot (M = −.0057, SEM = .0020, z = −2.83, p < .005, d = −.96) and marginally larger than object trials (M = −.0043, SEM = .0018, z = −2.36, p < .02, d = −.7). In the MOB, the rat-robot interactions showed significantly lower theta amplitude during grooming events (Int = .010, SEM = .002) than rat-rat interactions (M = .0028, SEM = .0009, z = 2.75, p < .003, d = .82) but no difference from objects (M = .0008, SEM = .0013, z = .65, d = .24). Rearing events during rat-rat interactions (Int. = .0015, SEM = .0001) exhibited significantly larger amygdalar theta oscillations than rearing events during rat-robot interactions (M = −.0002, SEM = .0001, z = −2.76, p < .006, d = −.59) but showed no difference from rat-object interactions (M = −7.2e-5, SEM = .0001, z = −.62, d = −.29).

#### 3.4.2 Respiratory Rhythms: Autonomic State and Distress Regulation

As an indicator of autonomic state and distress regulation, we predicted higher amplitude respiratory oscillations for the object and robot interactions during grooming (See Figure 8). Immobility events showed a significantly higher respiratory amplitude in the amygdala during the rat-robot interactions (Int. = .0032, SEM = .0001) than rat-rat interactions (M = −.001, SEM = .0003, z = −3.06, p < .0022, d = −1), but showed only a trend towards a significant difference between rat-object interactions (M = −.0004, SEM = .0002, z = −2.07, p < .04, d = −.4). Respiratory rhythm amplitudes in the amygdala during grooming events from rat-robot interactions (Int. = .0039, SEM = .0004) were significantly larger than events from rat-rat interactions (M = −.00016, SEM = .0006, z = −2.75, p = .006, d = −.8), but showed no significant difference from rat-object interactions (M = e-5, SEM = .0007, z = −.04, p = .96, d = −1.47e-2). Taken together these findings indicate that both robots and novel objects elicited an increase in arousal relative to conspecifics. Consistent with this interpretation, rearing also showed a significantly larger amygdalar respiratory amplitude for rat-robot conditions (Int. = .0022, SEM = .0002) when compared with rat-rat (M = −.0007, SEM = .0003, z = −2.55,p < .01,d = −.7), but not significantly different from rearing events from rat-object interactions (M = −.0001, SEM = .0003, z = .43, p = .67, d =-.1).

#### 3.4.3 Theta and High Beta Oscillations: Sensorimotor Exploration and Recognition

Although we did not have explicit prior hypotheses regarding the hippocampus and amygdala theta oscillations during grooming, we did expect that there may be differences revealing whether the rat was recognizing the robot more as an object or more as a conspecific (See Figure 8). Hippocampal theta amplitude during grooming events from rat-rat interactions were significantly different from the object conditions (M = −.0015, SEM = .0005, z = −3.05, p < .002, d = −1.5), but showed no difference when compared with robot (M = −.0009, SEM = .0001, z = −1.42, p = .16, d = −.9). The amplitude of theta oscillations in the amygdala during grooming events from rat-rat interactions (Int = .0013, SEM = .0001) were significantly larger than object conditions (M = −.0002, SEM = .0001, z = −2.58, p < .01, d = −1) but not significantly different from robot conditions (M = −.0001, SEM = .0001, z=1.28, p = .2, d = −.5). One implication of this is that movement dynamics of another rat may elicit larger theta than the moving robot or stationary objects. Although this may be obligatory movement coding in the hippocampus, this dynamic might also serve to monotonically index rat, robot, and object based on this parameter.

As observed in previous work, it was expected that the object would result in a robust burst at beta frequency in the hippocampus due to learning of the object stimulus (Rangel et al, 2015). Instead, the presence of increases in high amplitude beta were more indicative of high beta/low gamma activity observed in the hippocampus during object place associations (Trimper et al, 2017). The amplitude of beta during rearing events from rat-object interactions (Int. = .0005, SEM = e-5) were significantly higher than events from rat-rat (M = −.0002, SEM = 1e-5, z = −4.9, p < 1e-6, d = −2) and rat-robot interactions (M = −.0001, SEM = 1e-5, z −4.17, p < 1e-4, d = −1). This too indicates that the rat is likely associating the object with its place and differentiates the object from the robot and rat, both of which are moving.

## 4 Discussion

Animals are inextricably embedded within an environment and situated within a rich social world, bound to active exploration in recursive loops of perception and action (Kirsh & Maglio, 1995; De Jaegher & Froese, 2008). Brain dynamics rapidly and transiently switch from exploring the external world to evaluating the internal effect of the world on the organism (Marshall et al, 2017). Artificial systems often lack the complexity, adaptivity and responsiveness of animate systems, but can mimic kinematic and dynamic properties that inanimate objects often lack. It is an open question in the literature as to whether the rat perceives the robot as an animate or inanimate object. This study presents data to suggest that brain regions preferentially dissociate between rat, robot, and object based on sensorimotor exploration, salience, and autonomic distress regulation, indicating that the robot bears similarity to both a rat and an object. Primarily, this study outlines a general approach for such experiments that emphasize naturalistic interactions and complementary analysis pipelines that are necessary to render a holistic picture of the behavioral and brain dynamics evoked during interactive neurorobotics experiments.

Dynamic behavioral data regarding inter-agent distance suggest that the rat may be initially engaging in risk assessment behaviors when interacting with the robot, whereas they more readily approach another conspecific or stationary object. This may indicate that the rat feels safer approaching conspecifics and stationary objects than a robot. The distance between agents was also affected by the robot more often occupying the middle of the field and the rat showing a preference for the wall, a thigmotaxic strategy generally attributed to safety seeking (Lipkind et al, 2004). While that may have influenced the distance, the trajectories over time show a spiraling between rat and robot which suggests that the rat was actively avoiding the robot. The trajectories also suggest that as time passes the rat may become habituated to the robot and allow for decreased inter-agent distance at the end of a trial (see Figure 5). This differs from two prior robot-rat studies, the Waseda rat and the Robogator, where rats’ engagement with the robot is primarily enemy avoidance (Choi and Kim, 2010; Shi et al, 2013). Instead the rats in our study begin to engage in closer interactions across time rather than keeping their distance. This is more consistent with a behavioral study of the robot e-Puck and rats in which the robot elicits social behaviors from the rats (del Angel Ortiz et al, 2016).

In the present study, however, rats do show distress responses to the robot that exceed those to a conspecific or a novel object. However, they demonstrate maximal anxiety on the empty open field (used to test anxiety, Prut & Belzung, 2003); this indicates that the robot on the open field might provide comfort or that it presents a degree of behavioral competition between curiosity and anxiety. Specifically, interactions with a robot elicited or perturbed immobility, grooming, and rearing behaviors. The authors note that the rat-rat interactions resulted in few instances of immobility due to their active engagement, and reduced self-grooming likely due to the availability of social grooming by the other conspecific and the absence of distress with the conspecific. Rat conspecifics interacted, heavily engaging in coordinated exploration, following, interactive play, and anogenital exploration (also see Figure 5). This is consistent with findings detailed in a prior study (del Angel Ortiz et al, 2016).

From a design perspective, roboticists look for cues regarding whether the rat perceives the robot as more of an object or as an animal. Neuroimaging studies of human subjects viewing social androids and their movements addressed this question, discovering that the brain dissociates between androids and humans according to form and motion interactions (Saygin et al, 2012). Neural data in the present study demonstrate both robust and subtle differences to the rat, the robot, and the object. Specifically, hippocampal theta during grooming differentiates the rat from the object, however the robot (falling in between the two) is not significantly different from either. Here it is possible that the rat is coded according to its spatiotemporal dynamics, that are missing from the object, and that the moving robot carries features of each.

In contrast, theta dynamics in the MOB are similar for robot and object while differentiating the rat. This was expected based on the complexity of the inherent biological odors of the rat. The increased amplitude in respiratory rhythms during alert immobility in the amygdala to robot and object suggest autonomic regulation related to increased distress related arousal when compared with a rat. This suggests that interacting with a rat may be more engaging and less distressing. This finding is consistent with the behavioral interaction data. Thus, the brain appears to differentiate between a rat and our robots, but also distinguishes between the robot and objects. These findings provide support for the context-based modulation of brain signals (Kalueff et al, 2016). They also imply that through iterative design of robots, one could eventually produce a robot for which many regions of the brain do not readily distinguish the robot from a rat.

Future directions will also involve collecting a larger data set for the purpose of examining transient gamma activity, especially during investigation of the other agent or object. Rhythms like high gamma indicate active processing of the external sensory world, while low gamma is likely related to regulating an animal’s internal interoceptive milieu (Kay et al, 2009). It is recommended to use techniques such as burst detection to capture the more transient agent-based brain responses within behavior. Future approaches should also examine inter-regional communication, such as coherence and dynamic coupling (Fries, 2015; Breston et al, 2021). Additionally, future studies should also incorporate a richer repertoire of stimuli and robots.

Comparing the neurobehavioral states evoked by conspecifics, robots and objects may clue us into some of the minimal requirements for an animal to perceive artificial agents as social others. A key insight from Datteri (2020) about the philosophical foundations of the field is that interactive biorobotics experiments by themselves do not necessarily tell us about how organisms interact with conspecifics or predators, and suggests we should examine these interactions in their own terms before drawing unwarranted conclusions from the observations. This does not preclude a comparative approach, it just requires that we first take the robot case on its own terms and then compare it with data from social, object, and predator interactions.

Thus, this initial data from our exemplar rodent-centered design study suggest that agent-based comparisons within behavior are a promising direction moving forward in the field of interactive neurorobotics. A limitation of this study is that the robots’ motion dynamics were animated by human drivers, this introduces potential issues with anthropomorphism of interaction which can be dealt with better using automatic and autonomous robots. However, WoZ is a necessary step in the development of autonomous systems allowing for a diverse collection of movement dynamics that autonomous robots currently lack the complexity and sensitivity to exhibit. Follow up studies will be performed using autonomous robotic systems, such as PiRat (Heath et al, 2018). Autonomous robots allow for the manipulation of dynamics or functions that can be systematically manipulated according to model-based reasoning.

The results of this study also have implications for engineering design. From an engineering perspective, key considerations of social robots are safety, interactivity, and robustness (See SI Design Principles: Lessons Learned). The considerations must not only be viewed from the perspective of sound engineering design, but they must also be considered from the internal perspective of the animal meant to interact with the robot. For example, the observational data and quantitative analyses suggest that a key aspect of designing interactive robots is safety. In this case safety must be considered not only by making sure that the robot is designed with physical safety in mind but more importantly whether the animal interlocutor feels safe when interacting with the robot. Safety is a key first step towards the eventual goal of getting the animal to accept the robot as a potential social companion and, thus, a primary consideration in interactivity as well.

Safety and robustness must also be considered from the perspective of species specific behaviors. For rats, robot shells or covers should be designed so that it can be easily removed by users, but not by the animals, and made of a relatively non-porous material that can be easily cleaned in case of contamination by any substance. Robotic platforms made for interaction with non-human animals must be mostly water-proof. This is because urination and urine marking are common occurrences based on our observations, as they are natural features of interactivity, and thus the robot’s shell or exterior coating must effectively seal the electrical components from an animal’s urine. We observed multiple occasions where animal’s urine marked the field, stationary objects and occasionally other conspecifics. Urine marking is a communicative act, which can denote territoriality, dominance, and is full of rich social information (Leonardis et al, 2021). Marking may bias future interactions if the robot is contaminated with social odor from another conspecific.

Interactive robots have visual, auditory, olfactory, tactile and even gustatory aspects that should be actively taken into account during the design process. For instance, audition plays a key role. It is critical that interactive robots exhibit audio frequencies outside of the range of rodent distress calls, which induce panic, irritation, or stress responses. When designing interactive robots for animals it is also critical to take the animals preferred sensory systems into account, which for rats brings olfaction to the foreground and is why the robots in this study were tagged with olfactory stimuli. Contestabile et al (2021) used a dynamic moving object as a control for complex social stimuli and found that mice prefer complex social stimuli where more multisensory information such as tactile, visual, olfactory and auditory cues are available. They demonstrate that the object imbued with social odor does not recapitulate the complexities of social interaction, but instead the authors emphasize the importance of multisensory integration. Future approaches for designing interactions should emphasize the multisensory nature of the design problem.

Robotics has significant potential to offer animal research because its use enables experimenters to control complex experimental parameters and to test embodied computational models that interact with the real world (Webb, 2000). The dynamics of sociality are non-trivial and require convergent data regarding the model system. The holistic framework of capturing naturalistic behaviors in multiple contexts with fine-grained analyses, sets forth rich neural and behavioral data to scaffold the design process of future social robots (See SI Proposed Framework). The field of interactive neurorobotics allows for the examination of how robots evoke emergent behavior and brain dynamics in living creatures during a variety of agent-based interactions. This approach can be generalized to other animals and to humans. Social robotic interactions with humans also have affective dimensions that are indexed by autonomic signals and design principles can be improved by taking detailed behavioral interactions and neural signaling into account. It is essential that we also look beyond these narrow experimental contexts and generalize these lessons to our own technologically enmeshed world. While there is no putting out the fire of our increasing integration with autonomous systems, we have the opportunity to use the methods and insights gained from interactive neurorobotics to mold them into cooperative companions.

## Data Availability Statement

The raw data supporting the conclusions of this article will be made available by the authors, without undue reservation.

## Ethics Statement

All experiments and maintenance procedures were performed in an American Association for Accreditation of Laboratory Animal Care (AAALAC) accredited facility in accordance with NIH and Institutional Animal Care and Use Committee (IACUC) ethical guidelines and preapproved by the IACUC committee.

## Author Contributions

EJL, LKQ, and AAC designed the study. EJL, LKQ, AAC, RLM, LSe, LSc collected the data. LKQ performed neurosurgeries. EJL, RK, RLM, JW, LBG, performed the behavioral data analysis. LB and EJL performed the neural data analysis. EJL, AAC, LB, RLM, LSe, wrote and revised the article. All authors contributed to the article.

## Funding

This work was funded by NSF BRAIN EAGER 1451221; NIMH R01 5R01MH110514-02; Kavli Institute for Brain and Mind

## Acknowledgements

We would like to acknowledge the help and input from Nicklas Boyer, Scott Heath, Carlos Ramirez-Brinez, Jt Taufatofua, Ola Olsson, Marcelo Aguilar-Rivera, Emmanuel Gygi, Estelita Leija, and many more.

## Notes

### Competing Interest Statement

The authors have declared no competing interest.

