## Supplemental Information for "Interactive Neurorobotics: Behavioral and Neural Dynamics of Agent Interactions"

**Behavioral Video Coding**

Video was coded for behavioral epochs using ChronoViz (Fouse, Weibel, Hutchins, Hollan, 2011), as well as ELAN 6.0. The following epochs were extracted during the experiments/trials, with each event having variable length. Rearing was coded when the rat lengthened their spine and extended their nose in order to better investigate the environment with their paws either braced against an object or wall, or held against the rat's chest when free-standing. Immobility was the final coded behavior, in which a rat remained motionless and alert, while interpreting incoming environmental stimuli. There were three agent subcategories present within the condition, labeled as robot, object, and rat, indicating who or what the subject interacted with during each trial. 1,212 behavioral events were identified in the video data. During trials with the robot agent, there were 75 counts of immobility, and 71 counts of rearing. During the trials with the rat agent, there were 15 counts of immobility, and 39 counts of rearing. The trials with the object, had 128 counts of immobility, and 62 counts of rearing. For the open field trial, there were 119 counts of immobility, and 575 counts of rearing. 128 baseline epochs were pulled from the open field data from epochs where the animal is neither rearing or immobile. When asked to label 100 randomly drawn clips of the behavioral epochs, agreement between two independent raters for these behaviors was high (Cohen's Kappa = .9).

**Neural Network Offline Tracking Training and Validation Results**

Position tracking for the rat and the robot was performed with U-Net convolutional neural network trained using the Social LEAP Estimates Animal Pose (SLEAP) tracking system (Pereira et al., 2020). Videos were recorded at 29.97 FPS with a frame size of 720x480 pixels. SLEAP has been shown to be robust to multi-animal tracking issues with intersecting parts, close interaction, and swaps. The bottom-up approach was utilized to compute probability maps known as partial affinity fields for each frame of the video. Skeletal landmarks were estimated by computing a part confidence map and fitting gaussians for each part. Each body part estimation is the peak of the fit gaussian. The rats' skeletal tracking points were defined according to the specification of Sturman et al. (2020), including the nose, head center, left ear, right ear, neck, left side, body center, right side, left hip, right hip, and tail base. A Kalman filter was used to address the temporal association problem of shifts between frames to maintain identity of the skeletons.

The iRat tracker performed the most accurately based on 477 labels. The BOE robot tracker was trained on 1573 labels. 1/5th of the labels were set aside as the test set, while the remaining labels were included in the training set. The multi-animal rat tracker was trained on 6,700 labeled instances of rats from 3380 video frames from single and multi-rat trials. For accuracy results on the multi-rat tracker and robot trackers see Figure S1. For training and validation loss results see Figure S1.

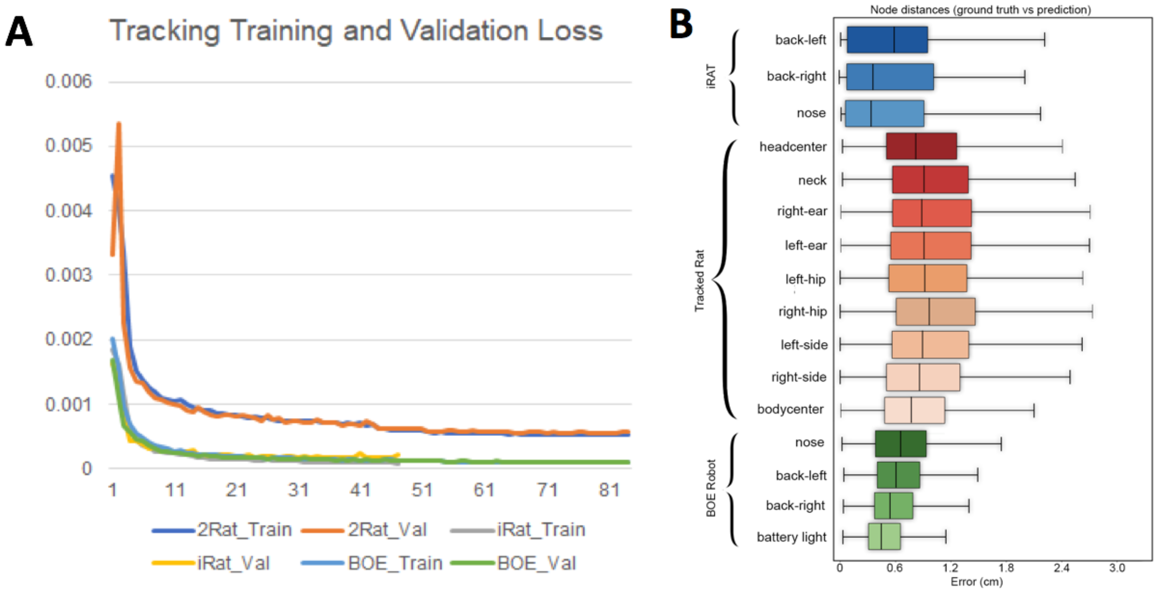


**Figure S1.** A, Training and validation results from neural network tracking of multi-rat, iRats and BOE robots. B, Accuracy results for the neural network tracker for multi-rat, iRat and BOE robots.

### Surgical Procedure

Rats (n = 3) underwent surgery for electrode implantation in order to record local field potentials from multiple brain areas simultaneously. Surgeries were performed in accordance with IACUC ethical guidelines. The rats were treated with isoflurane anesthesia (4-5% induction, 1-2% maintenance) and were placed in a stereotaxic apparatus to allow for placement localization (Kopf Instruments). Three holes were made through the skull, and the underlying dura was removed. LFP signals were referenced to a skull screw above the cerebellum. Anchor screws were inserted around the skull to support the neural implant which was cemented using dental cement. For the stereotrodes, pairs of 25 µm tungsten wire were twisted together and threaded through polyamide insulation. For the tetrodes, four sets of 12 µm wire were twisted together, and threaded through polyimide insulation (California Fine Wire). Electrodes were cut to the same length and wires were gold-plated in solution (Sifco) until impedances were reduced to approximately 100–300 kΩ measured at 1 kHz (Impedance tester IMP-1; Bak Electronics, Germantown, MD, USA). The stereotrodes were implanted using the stereotactic apparatus into the main olfactory bulb (8.5AP, 1.5ML, -3.5 DV), basomedial amygdala (-2.12AP, ±4.0ML, -9.2DV) and CA2/CA1 region of the hippocampus (-3.8AP, 3.8ML, 3.2DV) laterally.

### Neural Implants and Recordings

The stereotrodes and tetrodes were connected using gold pins to create contact between the wire and a Neuralynx E/I board cemented to the skull and anchor screws that send the electrical signal to an amplifier for signal processing. In the first rat, activity from 5 stereotrodes encased in polyamide tubing were connected to a 16-channel Neuralynx electrode interface board (EIB-16) that was cemented to the skull. The signals acquired from the E/I board were amplified using the Cheetah-32 system and Lynx-8 amplifiers (Neuralynx Technologies, Bozemon, MT). Amplifiers were integrated with the Cheetah data acquisition software provided by Neuralynx Technologies. The sampling rate for the recorded local field potentials was

1010.10Hz. Video was recorded from a camera above the field at 29.97FPS at 720x480 pixel resolution. Video was captured through the Cheetah data acquisition software, allowing for alignment between the timestamps of the neural data and video frames.

**Signal Processing**

Local field potential recordings from amygdala, hippocampus, and main olfactory bulb were indexed according to hand-coded behavioral epochs. To control for amplitude differences between subjects, LFP traces were normalized by overall standard deviation of the LFP per brain region for each rat. An infinite impulse response (IIR) bandstop filter was applied between 59-61Hz in order to filter out 60Hz line noise. Events with artifacts were detected using a .4 millivolt threshold on the CA1/CA2 and amygdala channels, and a .6 millivolt threshold on the MOB channels. An FIR bandpass filter was used to isolate the respiratory rhythm (2-6Hz), theta (6-10Hz), and beta (15Hz-35Hz). Although the respiratory rhythm commonly varies between 2 Hz and 12 Hz, the respiratory frequency overlaps with the theta so the range was restricted (Rojas-Libano et al, 2014). Shifts from respiratory to theta ranges often correspond to slower and faster sniffing rates (Tort et al, 2018). Power spectral densities and spectrograms were estimated using the Julia Fourier Analysis library, which is a windowed average across the log of the absolute value of the fast fourier transform (FFT) of the signal. The FFT decomposes the signal into frequency and amplitude features which can be used to identify the presence of oscillations or aperiodic rhythms in the signal.

**Wizard of Oz**

While ultimately the goal of interactive neurorobotics as a field will be to examine the interactions between rats and autonomous robots. This experiment is concerned primarily with how the form and motion of the robots compares with the animal data using Wizard of Oz methods and semi-autonomous robots to modulate the robots dynamics. The current experiment uses more traditional HRI methods, like WoZ and semi-autonomous methods. This experiment was performed using the iRat and DIY OpenSource Board of Education Shield robots. Robots and rats freely roamed the open field while the robot was controlled using Wizard of Oz mode (Riek, 2012). Experimenters controlling the robot were instructed to avoid dominance displays, cautiously approach, and modulate the position and movement dynamics to engage exploration and attentive behavior from the rat. The goal of the driver was to get the rat to follow, chase, or engage in tag-like play with the robot.

**Objects**

6 Objects (n = 6) were assembled by gloved hands with LEGO pieces. A small drop of diluted essential oils was placed on the back end of the object with a cotton swab so that they could more easily distinguish between the robots. Scents included frankincense, myrrh, tea tree, cedarwood, pine and rosemary. The objects had a length by width by height of 15x10x8cm. Much like the affordances of robots, the objects could support rearing and even climbing or mounting by the rat. Object position was varied from trial to trial to avoid place preferences.

**Board of Education Shield Arduino Robot (DIY Option)**

The robots were created using an Arduino UNO, an OpenSource prototyping microcontroller that executes code written in Arduino programming language, and the Board Of Education (BOE) Robot Shield Kit by Parallax Inc. These robots were used because they are readily available OpenSource robots that are easily accessible. The BOE kit includes the necessary parts for constructing a three wheeled robot with two continuous servo motors in the back of the robot, and one wheel on the front to allow for stability. Plastic Memorex CD cake boxes were melted and shaped to cover the electrical components of the device to ensure safety. This kit allows for remote-controlled driving by the experimenter (Wizard of Oz (WoZ) mode) or the execution of pre-programmed movements.

**Behavioral Hypothesis Tests:**

**Immobility:** Rats demonstrated few differences in immobility in all conditions except for the increased duration of immobility during the open field with no other rats, robots, or objects present. Mean frequency of immobility was marginally reduced between events from open field (M = 1.77, SEM = .15) when compared with rat-object interactions (M = 3.36, SEM = .72, t = -2.15, p < .03, d = .85). Mean frequency of immobility per trial was marginally larger during open field interactions than rat-rat interactions (M = 1.13, SEM = .3, t = 1.91, p < .04, d = .7). Mean duration of immobility events during interaction with the robot (M = 2.22, SEM = .09) show no significant difference from the rat (M = 1.95, SEM = .17, t = 1.47, p = .08, d = .24) and object interactions (M = 2.06, SEM = .11, t = 1.15, p = .13 , d = .14). The mean duration of immobility during open field interactions (M = 4.95, SEM = .30) was significantly longer than robot (t = 2.73, p < e-4, d =1.44), rat (t = 8.77, p < e-4, d = 1.15) and object (t = 9.06, p < e-4, d = 1.40) interactions.

**Grooming:** There were few differences in the frequency of grooming in the presence of rat, robot, or object. However, there is an increased duration for each groom demonstrated in the presence of the robot.There was no significant difference in the mean frequency of grooming events between the open field interactions and the rat-rat (M = 1.74, SEM = .22, t = .67, p = .25, d =.16) as well as rat-object interactions (M = 2.29, SEM = .53, t = .66, p = .25, d = .3). There was no significant difference between the mean duration of grooming between the open field interactions (M = 7.03, SEM = .65) and interactions with rat (t=.76, p > .05), robot (t=1.44, p = .08, d = .31), and object (t = -1.04, p > .05, d = .16).

**Rearing:** The mean frequency of rearing events during rat-robot interactions (M = 1.87, SEM = .22) showed no significant difference with rat-rat interactions (M = 1.54, SEM = .34, t = .80, p = .21, d = .25) or rat-object interactions (M = 2.58, SEM = .38, t = 1.62, p = .06, d = .49). The mean frequency of rearing during open field trials (M = 5.59, SEM = .35) was significantly larger than rat-robot (t = 8.9, p < e-4, d = 1.4), rat-object (t = 5.70, p < e-4, d = 1.04) and rat-rat interactions (t = 8.21, p < e-4, d = 1.4).

**Neural Hypothesis Testing**

There was no significant difference theta amplitude in MOB during grooming events between rat-robot (Int. .01, SEM = .002) and rat-object interactions (M = .0008, SEM = .001, z = .65, p = .5, d = .26)**.** Grooming events during rat-rat interactions exhibited no difference between the amplitude of the hippocampal theta oscillation (Int = .0054, SEM = .0007) when compared with robot (M = -.0009, SEM = .0001, z = -1.42, p = .16, d = -.9). There was no difference in MOB theta amplitude during rearing events from rat-rat interactions (Int. = .025, SEM = .006) when compared with the rat-robot (M = -.0008, SEM = .001, z = -.73, p = .46, d = -.18) and rat-object interactions (M = -.002, SEM = .002, z = -1.16, p = .25, d = -.38).

**Proposed Framework**

With its promise, interactive robotics brings a suite of challenges that must be addressed through a change of approach to experimental design. This shift is necessary because the standard doctrine of minimizing free variables and pursuing robust, repeatable effects, is in conflict with the method's primary raison d'etre– to capture the complexity of naturalistic behavior. What follows is a proposed framework, representing a synthesis of our lessons learned, for creating effective interactive robotic experiments that embrace its ability to capture the complexity of naturalistic behaviors while still allowing for robust scientific results. The framework can be summarized in terms of three goals:

1. Create as naturalistic behavior as possible within the experiment's objectives. Accepting the inherent complexity of agent interactions, instead of trying to constrain them, maximizes the method's utility because its primary strength is the ability to capture more naturally representative behavior. To accomplish this goal it is necessary to approach the design of both the robot and behavioral assay in terms of organism-centered design which prioritizes the comfort of the experimental subject. Furthermore, as it is advised to include conspecific and inanimate object controls to be able to estimate how the organisms robot interactions deviate from the natural target.
2. Collect comprehensive multimodal observations during the experimental trials. To make sense of the highly variable dynamics that occur during agent interactions it is necessary to record sufficient data to be able to reconstruct the behavioral and contextual subtleties that contribute to the variance. Additionally, combining multiple independent measurement modes provides a holistic perspective of the experimental results and can combine synergistically when building evidence for scientific conclusions. For measurement modes with constrained capacity, be careful to allocate those resources to components of the system with strong support for their involvement in the experimental assay.
3. Use a diversity of analyses to test a restricted number of hypotheses. The amount and variety of data, as well as the many degrees of freedom in the experiment itself present a danger of falling into a trap of making too many comparisons to report any results with certainty. This can be avoided by using many independent types of analysis to jointly support a small set of well supported hypotheses.

**Design Principles: Lessons Learned**

Three important features of our robotic design are safety, interactivity, and robustness. Safety is a critical aspect of autonomy, so significant measures must be taken to avoid any harm to the rodents such as running over or pinching their tail or bumping into them. Multiple protocols were created to avoid collision with the rat, the walls of the field, or any objects in the environment. Safety must also be factored in when designing the shape of the chassis and cover. Care should be taken to ensure that all edges are rounded so that both rats and users would not be harmed or intimidated by the robot. The cover should also completely contain all mechanical and electrical components, preventing any damage to those interacting with the robot as well as protecting the robot itself.

Robustness is crucial and can be increased overall by increasing the durability and strength of individual components (Kragic, 2004). Falling under both safety and robustness constraints, covers should be designed so that it can be easily removed by users, but not by the animals, and made of a relatively non-porous material that can be easily cleaned in case of contamination by any substance. Robotic platforms made for interaction with non-human animals must be generally water-proof. This is because urination and urine marking are common occurrences based on our observations, thus the robot's shell or exterior coating must effectively seal the electrical components from an animal's urine. We observed multiple occasions where animal's urine marked the field, stationary objects and occasionally other conspecifics. Cloth, fibers, or other such materials should be avoided or carefully tucked away from the rats' access to prevent choking, shredding, or scent marking that would damage either the rat or the device. A robot's weight should also be considered and be carefully weighted for the agents it will be interacting with. If the robot is too light, it can be easily tipped over, dented or crushed in. If the robot is too heavy, it can impact the motors, adding strain, reducing speed or maneuverability, or cause damage to the environment or to the rodents.

Interactivity consists of multiple sensory stimuli implemented to engage with the rats, with highly stimulating robots contributing to a more engaging environment. However, these stimuli must not be overwhelming, irritating or frightening. Rats can hear frequencies around 250 Hz to 80 kHz, able to hear ultrasound as well as being highly sensitive to frequencies between 8 kHz and 38 kHz (Escabi et al., 2019). It is critical that interactive robots exhibit audio frequencies outside of the range of rodent distress calls, which induce panic, irritation, or stress responses. Ultrasonic motors or traditional servos generate too many high frequency sound emissions, as we discovered during the piloting and recording of ultrasonics. The solution for the PiRat was to use gimbal motors which were sufficiently quiet within the necessary frequency range (Heath et al, 2018). Olfactory tags were used in this experiment to engage the rodent's dominant sensory system. Each of the two iRats were tagged with either frankincense or myrrh essential oils, meant to comply with preference profiles indicative of a naturalistic, woody scent.

While it may seem making the robot more like a rat will elicit naturalistic social behaviors, that is complicated by the possibility of the "uncanny valley" effect for animals. While this is primarily an effect associated with humans (Mori, 1970; Saygin et al, 2012), previous studies like those with the Waseda rat showed that the rat-like robot resulted in an anxiety response (Ishii et al, 2006; Shi et al, 2013). This would mean that the closer the robot is to having that animal's appearance the more of a distressing stimulus it may be due to bearing high similarity to a rat but having some features that are a mismatch.

The iRats are visually unobtrusive, with a tapered, rounded 'nose', and a wider backside. They are approximately three to four inches tall and four to five inches in length. In addition to its rounded shell, the cover of iRat is smooth, hard plastic, durable enough to support the weight of a rat without damage to the robot and without pricking the paws of the rat. iRat maneuvers the field with smooth, coordinated movements, pivoting around objects and rats at a sufficiently moderate speed so as not to provoke the rats, though the pace can be adjusted by the researchers so as to more accurately mimic the quick darting pace of a rat at play or the subdued crawl of a prowling robot. When associating the robot with a reward it is highly recommended that the robot and the reward site be separated from the agent. Gianelli et al. (2018) and Waseda Mouse No.8 had the robot itself act as the pellet dispenser. Ishi et al (2006) has shown separately that robots can easily teach rats to receive a reward at a nearby pellet dispenser. Interactive neurorobotics platforms would benefit from segregating the reward in order to avoid the confounding of extrinsic reward learning in the brain with the intrinsic reinforcement of the robotic stimulus.
